# Indole Pulse Signalling Regulates the Cytoplasmic pH of *E. coli* in a Memory-Like Manner

**DOI:** 10.1101/425538

**Authors:** Ashraf Zarkan, Santiago Caño Muñiz, Jinbo Zhu, Kareem Al Nahas, Jehangir Cama, Ulrich F. Keyser, David K. Summers

## Abstract

Bacterial cells are critically dependent upon pH regulation. Most proteins function over a limited pH range and the pH gradient across the bacterial cell membrane is central to energy production and transduction^1^. Here we demonstrate that indole plays a critical role in the regulation of the cytoplasmic pH of *E. coli*. Indole is an aromatic molecule with diverse signalling roles that in bacteria is produced from tryptophan by the enzyme tryptophanase (TnaA)^2^. Two modes of indole signalling have been described: persistent and pulse signalling. The latter is illustrated by the brief but intense elevation of intracellular indole during stationary phase entry^3,4^. We show that *E. coli* cells growing under conditions where no indole is produced maintain their cytoplasmic pH at 7.8 ± 0.2. In contrast, under conditions permitting indole production, pH is maintained at 7.2 ± 0.2. Experiments where indole was added experimentally to non-producing cultures showed that pH regulation results from pulse, rather than persistent, indole signalling. Furthermore, the application of an artificial pulse of either of two non-biological proton ionophores (DNP or CCCP) caused a similar effect, suggesting that the relevant property of indole in this context is its ability to conduct protons across the cytoplasmic membrane^5^. Additionally, we show that the effect of the indole pulse that occurs normally during stationary phase entry in rich medium remains as a “memory” to maintain the correct cytoplasmic pH until entry into the next stationary phase. The indole-mediated reduction in cytoplasmic pH may explain why indole provides *E. coli* with a degree of protection against stresses, including some bactericidal antibiotics.

## INTRODUCTION

Bacteria have evolved to grow over a wide range of external pH, while maintaining their cytoplasmic pH within a narrow range; a phenomenon known as pH homeostasis^6^. *E. coil* is a neutrophilic bacterium that can grow at external pH values from 5.0 to 9.0 but generally maintains its cytoplasmic pH in the range 7.2-7.8^1,8^. Recently, intracellular alkalisation has been proposed as a mechanism of bacterial killing by antibiotics^7^, making pH control a potentially important mechanism for countering the bactericidal activity of these drugs^1,8^.

Indole is an aromatic molecule produced by over 85 species of bacteria with multiple and diverse signalling roles^2^. Indole signalling in bacteria can be divided into two main types: persistent signalling and pulse signalling. In persistent signalling, the indole signal is present in the culture supernatant for an extended period at a relatively low concentration (<1 mM) and under these conditions it has been shown to regulate biofilm formation^9,10^, virulence^11^ and response to various stresses including antibiotics^12^. In pulse signalling, intracellular indole is transiently at a much higher concentration than in the culture supernatant. For example, in *E. coli* during stationary phase entry the cell-associated concentration reaches 50-60 mM for 10-20 min. Indole is a proton ionophore and the pulse is proposed to reduce the electrical potential across the cytoplasmic membrane, inhibiting growth and cell division^5^. This promotes the transition from exponential to stationary phase before nutrients are completely exhausted, with benefits for long-term survival^3,4^.

Although pH has been proposed previously to control indole production in *E. coli*^2^, the effect of indole signalling on pH regulation remains unexplored. Such a study requires the measurement of cytoplasmic pH that has previously been attempted by a range of techniques, including phosphorus nuclear magnetic resonance (^31^P NMR)^13^ radiolabelled transmembrane probes^14^ and pH-sensitive fluorescent proteins^15,16^. The latter seems to provide the best platform for rapid and highly-sensitive detection of bacterial cytoplasmic pH^17^. In the present study, we tracked the cytoplasmic pH of *E. coli* growing in LB and M9 media using a plasmid-encoded pH-sensitive fluorescent protein, pHlourin^15,16^. The cytoplasmic pH was maintained at 7.2 ± 0.2 in conditions that allowed indole production but switched to 7.8 ± 0.2 in its absence. An experimentally-applied indole pulse restored the pH of indole non-producing cells to 7.2. A similar result was achieved by a pulse with synthetic proton ionophores, suggesting that an ionophore mechanism is responsible for pH regulation. We have also shown that the cytoplasmic pH in exponential phase cells is set by the indole pulse which took place during the preceding stationary phase entry. This implies the presence of a cellular “memory” that can maintain the cytoplasmic pH for many hours after the pulse has subsided. We speculate that indole-mediated pH regulation may have important consequences for bacterial stress responses, including their ability to survive antibiotic treatment.

## RESULTS

### The difference in cytoplasmic pH between *E. coli* grown in minimal and rich media is due to indole production

Cytoplasmic pH was tracked using an mCherry-pHluorin translational fusion protein expressed from a pBAD TOPO vector (Invitrogen) under arabinose induction. pHluorin (also known as Super-ecliptic pHluorin) is a pH-sensitive green fluorescent protein^15,16^. mCherry fluorescence is not affected by pH and was therefore used to normalise changes in fluorescence caused by cell-to-cell variation in protein expression and differences in plasmid copy number between exponential and stationary phase. Using the mCherry-pHluorin plasmid, the cytoplasmic pH of *E. coli* BW25113 wild-type (WT) was monitored in LB or M9/glucose (0.4%). Overnight cultures were diluted to OD_600_ =0.05 in fresh LB or M9 medium at 37 °C and sampled at intervals between OD_600_ = 0.1 - 4.0 for M9 and OD_600_ = 0.1 – 6.0 for LB (Supplementary Figures; Fig. S1).

The cytoplasmic pH of *E. coli* BW25113 wild-type remained constant throughout the growth experiment but the absolute value was medium-dependent: 7.2 ± 0.2 in LB and 7.8 ± 0.2 in M9/glucose (Fig. 1a). To test whether the difference was due to some component present in the complex (LB) medium but absent in M9/glucose, trace elements, vitB1 and casamino acids were added to M9 (supplemented M9) but the cytoplasmic pH remained unchanged at 7.8. Casamino acids are obtained by acid hydrolysis of casein which destroys tryptophan^18,19,20^. Thus, tryptophan (1 mM) was added separately to supplemented M9 to provide a complete set of amino acids. In this tryptophan-containing defined medium the cytoplasmic pH of *E. coli* BW25113 became the same as in LB (Fig. 1a).

**Fig 1:**
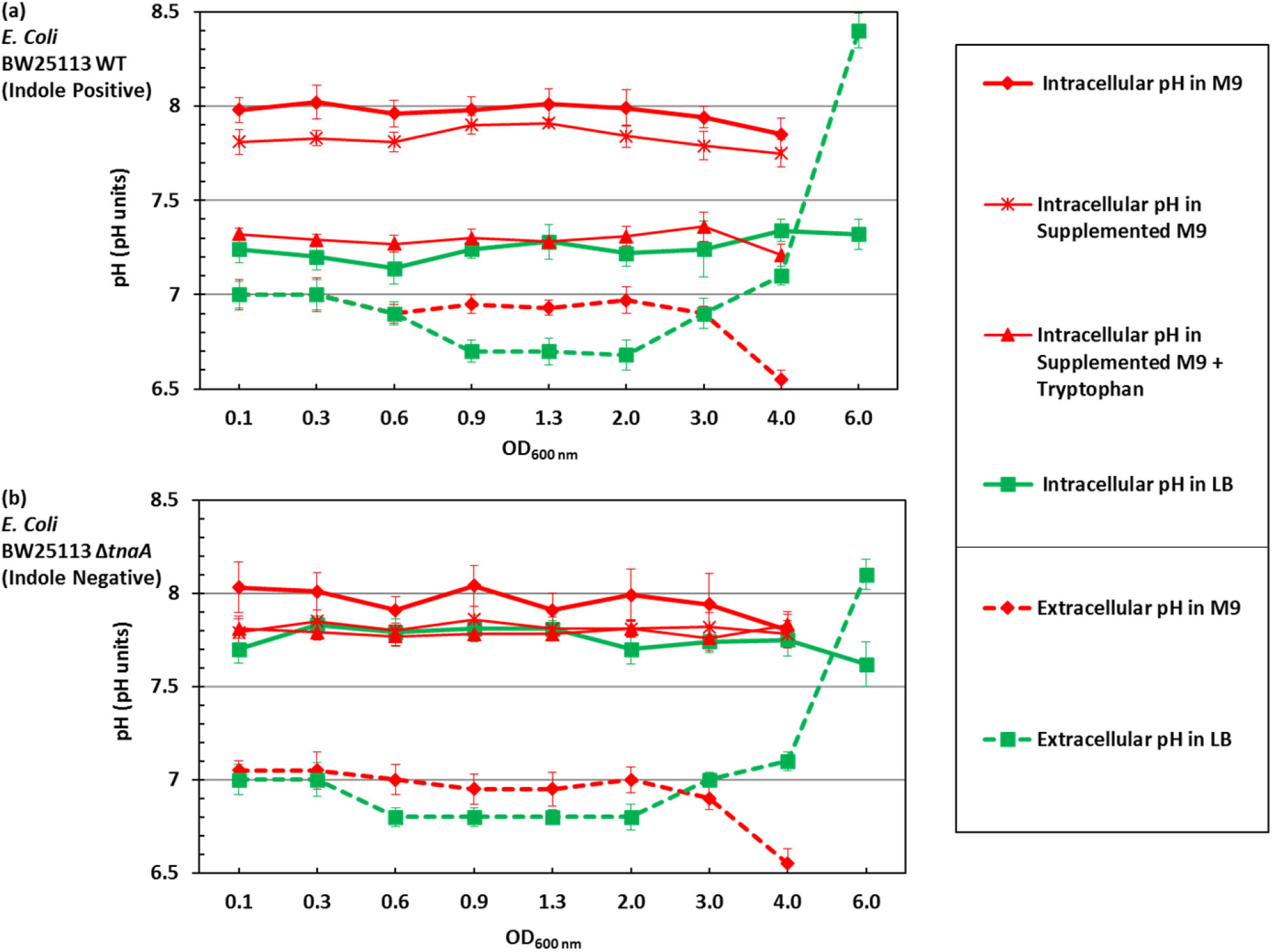
External and internal pH of *E. coli* growing in LB and M9/glucose. (a) The cytoplasmic pH of *E. coli* BW25113 WT was monitored throughout exponential and stationary phase in LB, M9/glucose, supplemented M9/glucose (containing trace elements, VitB1 and casamino acids) and supplemented M9/glucose with added tryptophan. The extracellular pH was also measured in LB and M9/glucose. (b) The cytoplasmic and extracellular pH of the indole-negative mutant *E. coli* BW25113 ∆*tnaA* was measured in the same range of media described in (a). Data are presented as the mean ± SD of three biological replicates, and the final point in each line is an overnight sample (~ 18 h incubation).

The role of tryptophan in modulating cytoplasmic pH may be direct or indirect. Since tryptophan is converted by tryptophanase (TnaA) into the signalling molecule indole, the experiment was repeated in a ∆*tnaA* mutant strain, which is incapable of indole production. Tryptophan uptake by *E. coli* is *via* tryptophan permease (TnaB), which is still produced in the ∆*tnaA* mutant so the indole-negative strain remains capable of internalising tryptophan even though it cannot convert it to indole. In the ∆*tnaA* strain (Fig. 1b), the cytoplasmic pH of cells grown in LB, M9/glucose, supplemented M9 or supplemented M9 with tryptophan was around 7.8. Thus, indole, not tryptophan, is primarily responsible for the lowering the cytoplasmic pH, and this was confirmed by directly assaying each culture for indole and showing that only indole-producing cultures had a cytoplasmic pH of 7.2 (Supplementary Figures; Fig. S2). The pH values of the culture media were also measured. The pH of LB remained between 6.7 - 7.0 throughout exponential and early stationary phase but rose to over 8.0 in late stationary phase (overnight incubation). M9/Glucose remained around 7.0 throughout exponential and early stationary phase but dropped to around 6.5 in late stationary phase (overnight incubation). The results were similar for both the WT (Fig. 1a) and the ∆*tnaA* (Fig. 1b) strains. The pH rise in LB has previously been linked to the toxification of the medium^21,22^ while the drop in M9/glucose is probably due to the presence of glucose, as growth on glucose is known to lower the pH of the medium^22^.

### Validation of cytoplasmic pH measurements by flow cytometry

The measurements of cytoplasmic pH shown in Fig. 1 were performed by fluorescence spectroscopy and represent a population average of a large number of cells. Such an approach is potentially misleading if, for example, the culture consists of sub-populations of cells with different cytoplasmic pH. It is also possible that fluorescent protein, released into the growth medium by cell lysis, could affect the result. To test the validity of the fluorimeter technique and to explore the possibility of sub-populations with different cytoplasmic pH, cultures were analysed by flow cytometry under two laser illuminations: blue (Ex 488 nm) for detecting the fluorescence of pHluorin and yellow (Ex 561 nm) for detecting the fluorescence of mCherry. The fluorescence distributions of pHluorin and mCherry show single peaks for both the WT (Fig. 2a) and the ∆*tnaA* mutant samples (Fig. 2b) suggesting relative uniformity of cytoplasmic pH among cells within each culture. The average intensity of the pHluorin and mCherry signal increased after entry into stationary phase (OD_600_ ≥ 2.0), probably reflecting plasmid amplification and confirming the need to normalise the pHlourin signal. The assertion that cytoplasmic pH has an effect on the fluorescence of pHluorin, but not mCherry, was confirmed by changing the external pH (6.5, 7.0, 7.5 and 8.0) of samples from both cultures (OD_600_ = 0.6) and treating them with carbonyl cyanide *m*-chlorophenyl hydrazone (CCCP) to equilibrate the cytoplasmic and external pH. The average intensity of pHluorin increased with increasing the pH, while mCherry remained unaffected. When these data were used to calculate average cytoplasmic pH for the wild-type and ∆*tnaA* cultures, the results closely reproduced those obtained by fluorescence spectroscopy (Supplementary Figures; Fig. S3).

**Fig 2:**
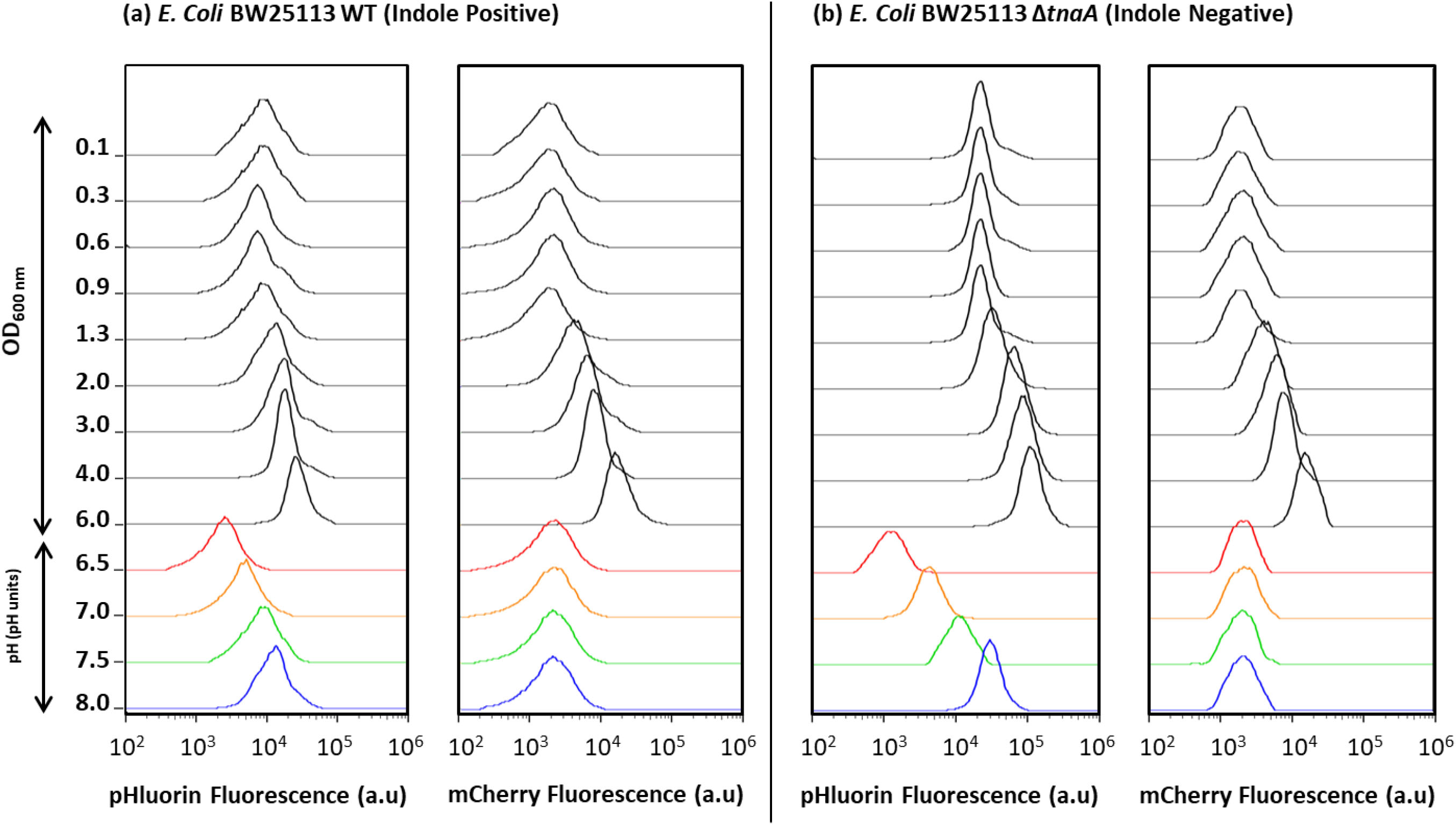
Distributions of single cell fluorescence (pHluroin and mCherry) used to calculate cytoplasmic pH. Samples from LB cultures of (a) BW25113 WT and (b) BW25113 *ΔtnaA* were analysed by flow cytometry. For each strain the fluorescence distributions of pHluorin and mCherry are shown for cultures initially growing exponentially and then entering stationary phase. The final sample in both strains (OD_600_ = 6.0) was taken after overnight incubation (~ 18 h incubation). Culture densities for each sample are shown in the Y-axis. The bottom four traces in each column are samples taken from the cultures at OD_600_=0.6 and exposed to a range of external pH in the presence of CCCP (which equilibrates the external and cytoplasmic pH).

### Indole pulse signalling maintains the cytoplasmic pH of *E. coli* at pH 7.2

Our comparison of wild-type and ∆*tnaA E. coli* suggests that indole production is necessary to maintain the cytoplasmic pH of the wild-type at 7.2. This implies that it might be possible to “repair” the pH of the ∆*tnaA* mutant by adding indole to the culture. The effect of indole is exerted through one of two modes: persistent or pulse signalling. Persistent signalling occurs at the concentrations of indole found in the culture supernatant. Indole concentrations in stationary phase LB cultures are in the range 0.3–0.7 mM with most indole synthesised during stationary phase entry^5,23,24^. The concentration in exponential phase cultures is much lower but quantitative data are not available. To improve the accuracy of the measurement, indole in the culture supernatant during the lag and exponential phase was concentrated 10-fold using a C18 solid phase extraction cartridge prior to assay. The assay revealed that although indole was undetectable immediately after sub-culture, it rose rapidly to 30 ± 10 µM during lag phase and remained approximately constant throughout exponential phase (Supplementary Figures, Fig. S4).

To investigate whether indole persistent signalling was responsible for regulating the cytoplasmic pH of *E. coli*, cultures of BW25113 ∆*tnaA* in LB were supplemented with a range of indole concentrations from 30 µM to 0.5 mM, and the cytoplasmic pH was measured using the fluorimeter method. Irrespective of the concentration, there was no significant change in the cytoplasmic pH of the ∆*tnaA* mutant strain (Fig. 3a).

**Fig 3:**
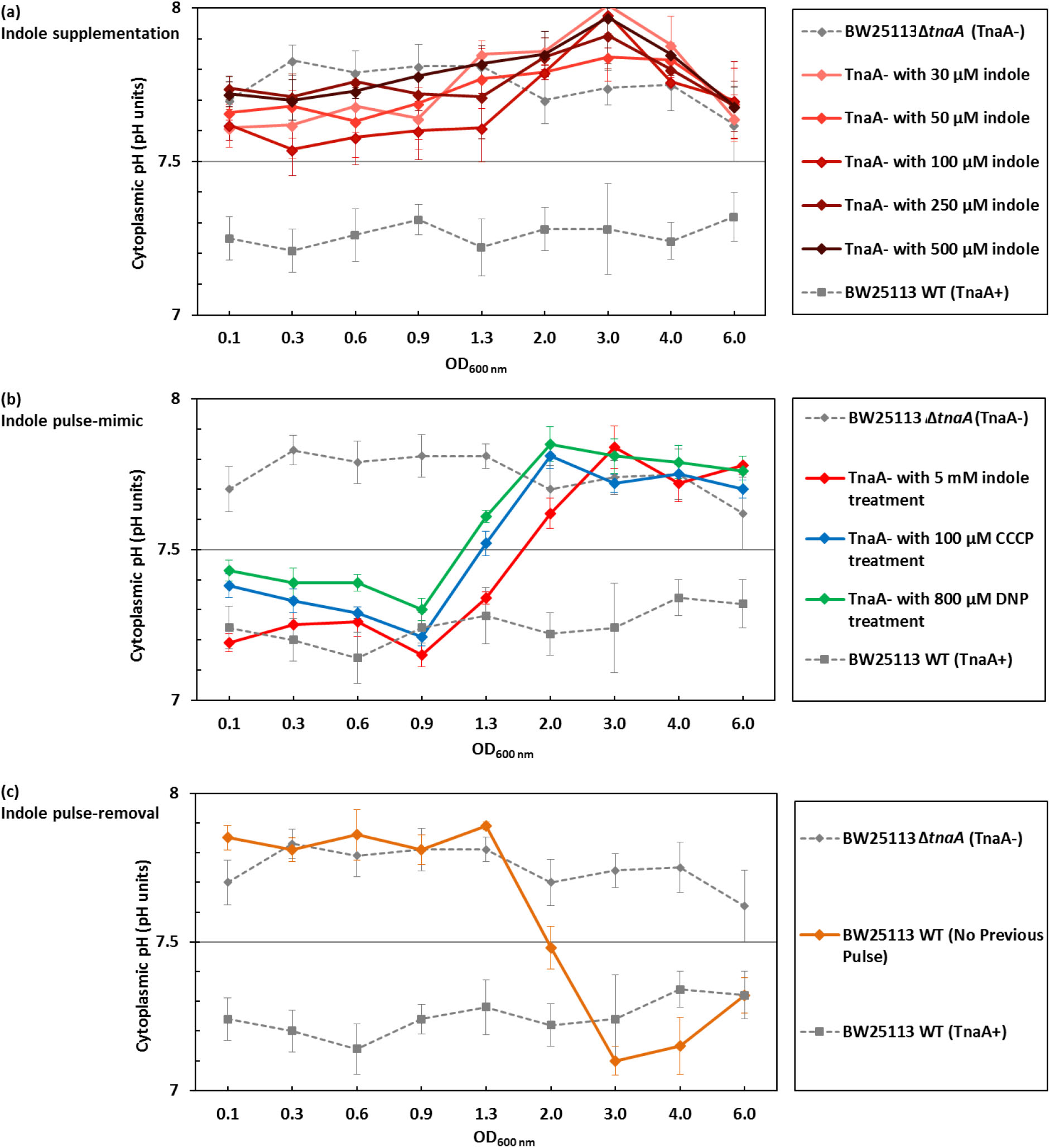
Indole pulse signalling regulates the cytoplasmic pH of *E. coli* by an ionophore-based mechanism. **(a)** To mimic the effect of indole persistent signalling, LB was supplemented with indole (30 - 500 µM) and the cytoplasmic pH of BW25113 **∆***tnaA* cells was monitored through exponential and stationary phase. **(b)** To mimic the effect of indole pulse signalling, BW25113 **∆***tnaA* cells in early exponential phase (OD_600_ = 0.1) were treated with 5 mM indole for 20 min. Cells were resuspended in LB (no indole) and their cytoplasmic pH was monitored through exponential and stationary phase. The result was compared with the effect of a 20 min pulse with non-biological ionophores, CCCP (100 µM) and DNP (800 µM). **(c)** A BW25113 WT culture was grown overnight (~ 18 hrs) in M9/glucose with no added tryptophan and sub-cultured the following day into LB (OD_600_ = 0.05). The cytoplasmic pH was monitored through exponential and stationary phase. Data for the cytoplasmic pH of BW25113 WT and ∆*tnaA* in LB presented in Fig. 1 were added to each of the three panels for comparison. All data are the means ± SD of three biological replicates, and the final point in each line was an overnight sample (~ 18 hrs incubation).

The effect of indole pulse signalling on cytoplasmic pH was investigated with an experimentally-applied indole pulse. The peak cell-associated indole concentration that occurs during the transition of wild-type cells from exponential to stationary phase is approximately 60 mM, and this can be mimicked experimentally by the addition of 4-5 mM indole to the culture supernatant^3^ (the difference between the external and cell-associated concentrations is due to the greater affinity of indole for cells than for the aqueous culture medium^3^). We treated an early exponential phase LB culture of ∆*tnaA* cells with 5 mM indole for 20 min to determine the effect of an indole pulse. Afterwards, the indole was removed by resuspending the cells in fresh LB and the cytoplasmic pH was monitored during the remainder of exponential phase and into stationary phase. The pH of ∆*tnaA* cells in the culture that had been subjected to the artificial pulse was indistinguishable from that of a control wild-type culture (7.2 ± 0.2) until the transition from exponential to stationary phase at around OD_600_=1.0 (Fig. 3b). Thereafter the cytoplasmic pH reverted to 7.8 ± 0.2. It is of interest that reversion to the higher pH took place at the same time that the next indole pulse would naturally have occurred in a wild-type culture.

Having demonstrated that the wild-type pH could be restored in a ∆*tnaA* mutant strain by an experimentally-imposed indole pulse, we next investigated the effect of removing the pulse on the cytoplasmic pH of wild-type cells. We have reported previously that wild-type cells growing in medium lacking tryptophan do not produce an indole pulse during stationary phase entry^4^. Wild-type cells were cultured overnight (approx. 18 hrs) in M9/ glucose without added tryptophan and sub-cultured the following day into fresh LB. Thus, the cells had no pulse during the overnight culture but, once sub-cultured into tryptophan-containing medium, regained the ability to produce indole. As the cells sub-cultured in LB entered exponential phase, their cytoplasmic pH was indistinguishable from that of a ∆*tnaA* mutant strain (Fig. 3c). However, after the culture transitioned from exponential to stationary phase, with the accompanying indole pulse, the cytoplasmic pH changed to become indistinguishable from that of a wild-type culture.

### The indole pulse regulates the cytoplasmic pH of *E. coli* by an ionophore-based mechanism

At higher concentrations, indole acts as a proton ionophore^3,4,5^. If the regulation of cytoplasmic pH by indole involves a simple ionophore mechanism, it should be possible to replace indole with an alternative ionophore. When an artificially-imposed pulse of the non-biological ionophore CCCP (100 µM) or DNP (800 µM) was applied to an early exponential phase cultures of BW25113 *∆tnaA*, the cytoplasmic pH was restored to 7.2 ± 0.2 until the time of the transition to stationary phase when it reverted to 7.8 ± 0.2 (Fig. 3b). In other words, it responded to CCCP and DNP in exactly the same way as it responded to an experimentally-applied pulse of indole. Thus, we conclude that indole regulates cytoplasmic pH by an ionophore-based mechanism.

If indole changes cytoplasmic pH by allowing the flow of protons down a concentration gradient across the cytoplasmic membrane, the direction of the change should depend on whether the pH of the growth medium is higher or lower than the cytoplasm. Two trizma-buffered versions of LB were used to test our hypothesis; one at pH8 (Fig. 4a) and the other at pH7 (Fig. 4b). The cytoplasmic pH of the indole-deficient strain BW25113 ∆*tnaA* was 7.8 ± 0.2 in both media. However, the cytoplasmic pH of the wild-type strain was dependent on the external pH, shifting from 7.2 ± 0.2 in the pH7 medium to 7.6 ± 0.1 in the pH8 medium. Note that the medium lost its buffering capacity in late stationary phase, probably due to the accumulation of alkaline waste products.

**Fig 4:**
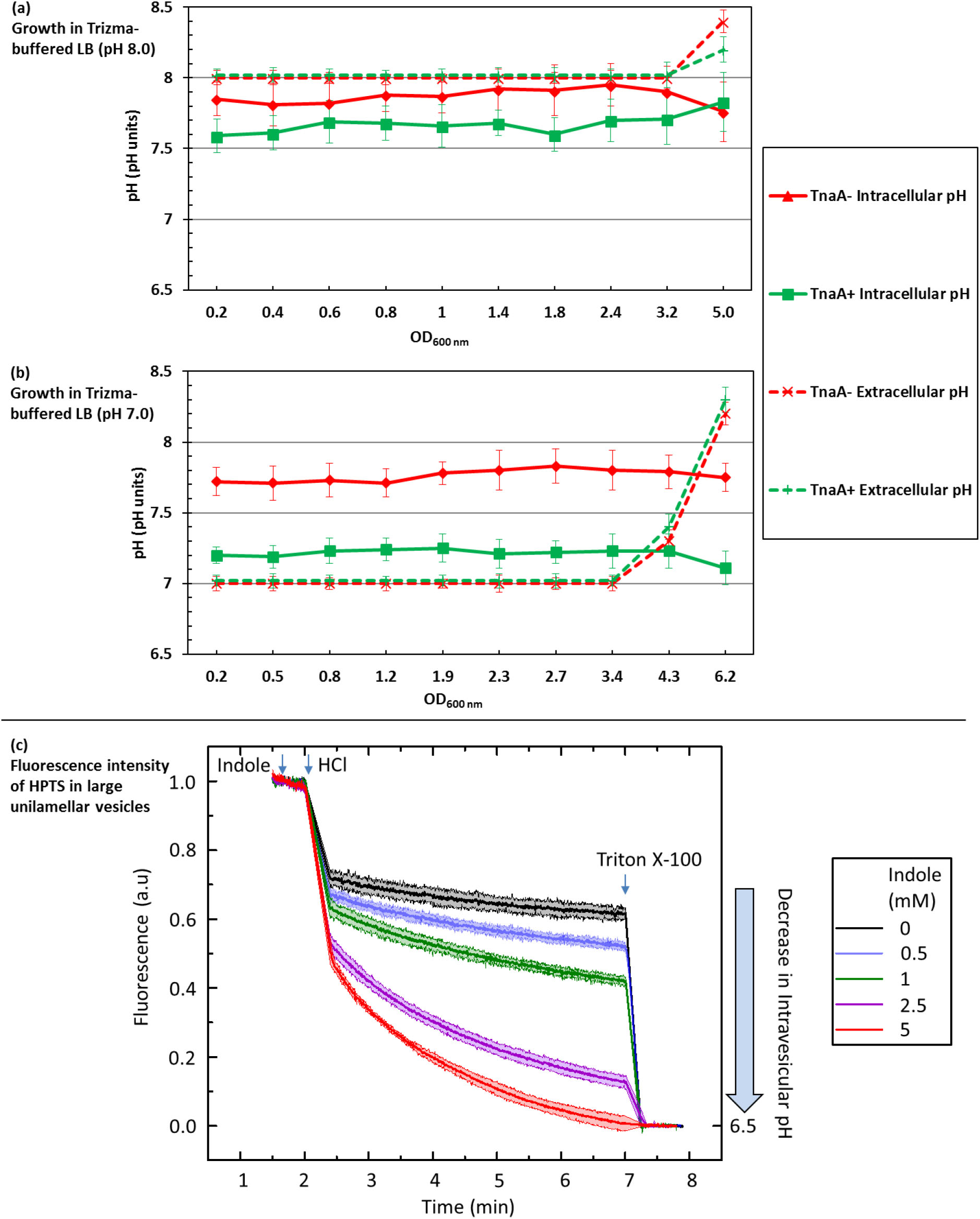
The relationship between external and internal pH of *E. coli* cells and large lipid vesicles. Comparison of the cytoplasmic pH of *E. coli* BW25113 WT and ∆*tnaA* growing in trizma-buffered LB at (a) pH 8,0 or (b) pH 7.0. Data are the mean ± SD of three biological replicates, and the final point in each line represents an overnight sample (~ 18 hrs incubation). (c) the fluorescence intensity of HPTS (*λ*_ex_=450 nm *λ*_em_=510 nm), an indicator of intravesicular pH, in large unilamellar vesicles (LUVs), was measured in the presence of indole (0-5 mM). Data presented are the mean ± SD of three biological replicates.

The buffered medium experiment (Fig 4a and b) suggests that the proton flux associated with an indole pulse is responsible for altering the pH of the cytoplasm. We used a model system of large unilamellar vesicles (LUVs)^25^ to quantify the concentration of indole required to achieve a significant pH change. We prepared LUVs encapsulating the pH-sensitive fluorescent dye, 8-hydroxypyrene-1,3,6-trisulfonic acid (HPTS)^26^ and the extravesicular and intravesicular pH were buffered at 7.4. Indole (0-5 mM) was added to the extravesicular buffer, followed by sufficient HCl to lower the external pH to 6.5. HPTS fluorescence (reflecting the internal vesicle pH) was monitored over the next 5 min, before the vesicles were lysed by adding the detergent Triton X-100, bringing HPTS rapidly to pH 6.5. The effect of indole was strongly dose-dependent. The addition of 5 mM indole caused the intravesicular pH to fall to 6.5 within 5 min. However, at 0.5 mM (typical of the indole concentration in LB stationary phase culture) there was only a small reduction in the intravesicular pH, compared to the no-indole control (Fig. 4c). Our result supports the hypothesis that an indole concentration characteristic of pulse signalling would be required to cause a significant change in the pH of the *E. coli* cytoplasm.

## DISCUSSION

Our experiments demonstrate the existence of an indole-mediated mechanism for the regulation of cytoplasmic pH in *E. coli*. We have shown that a lack of indole production, due either to deletion of the tryptophanase gene or by growing cells in medium lacking tryptophan, causes the cytoplasmic pH of *E. coli* to shift to 7.8 ± 0.2, compared to 7.2 ± 0.2 when indole is produced (Fig. 1).

The regulation of pH by indole seems to be entirely dependent upon pulse signalling during stationary phase entry. Supplementation of the growth medium of tryptophanase knock-out cultures with 30-500 µM indole (representative of concentrations found in the culture supernatant of exponential or stationary phase cultures) failed to restore the lower cytoplasmic pH (Fig. 3a). However, an artificial indole pulse rapidly restored the cytoplasmic pH of the tryptophanase knock-out strain to the wild-type value. When wild-type cells are growing in tryptophan-containing medium, the indole pulse which occurs during the transition from exponential to stationary phase is the key to indole-mediated pH regulation. When this pulse was prevented by growing an overnight culture of wild-type cells in the absence of tryptophan, the cytoplasmic pH of sub-cultured cells in fresh LB was similar to that of ∆*tnaA* rather than wild-type cells, even though the sub-cultured cells had access to tryptophan in LB and were genetically capable of indole production. The “mutant pH” was maintained until the subsequent transition to stationary phase when it was corrected by the associated indole pulse. Thus, the pulse is not only capable of setting the cytoplasmic pH of *E. coli* entering into stationary phase but also maintains it at the correct value during exponential phase growth following sub-culture into fresh medium the next day. This phenomenon, which we term the “pH cycle” is illustrated in Fig. 5.

**Fig 5:**
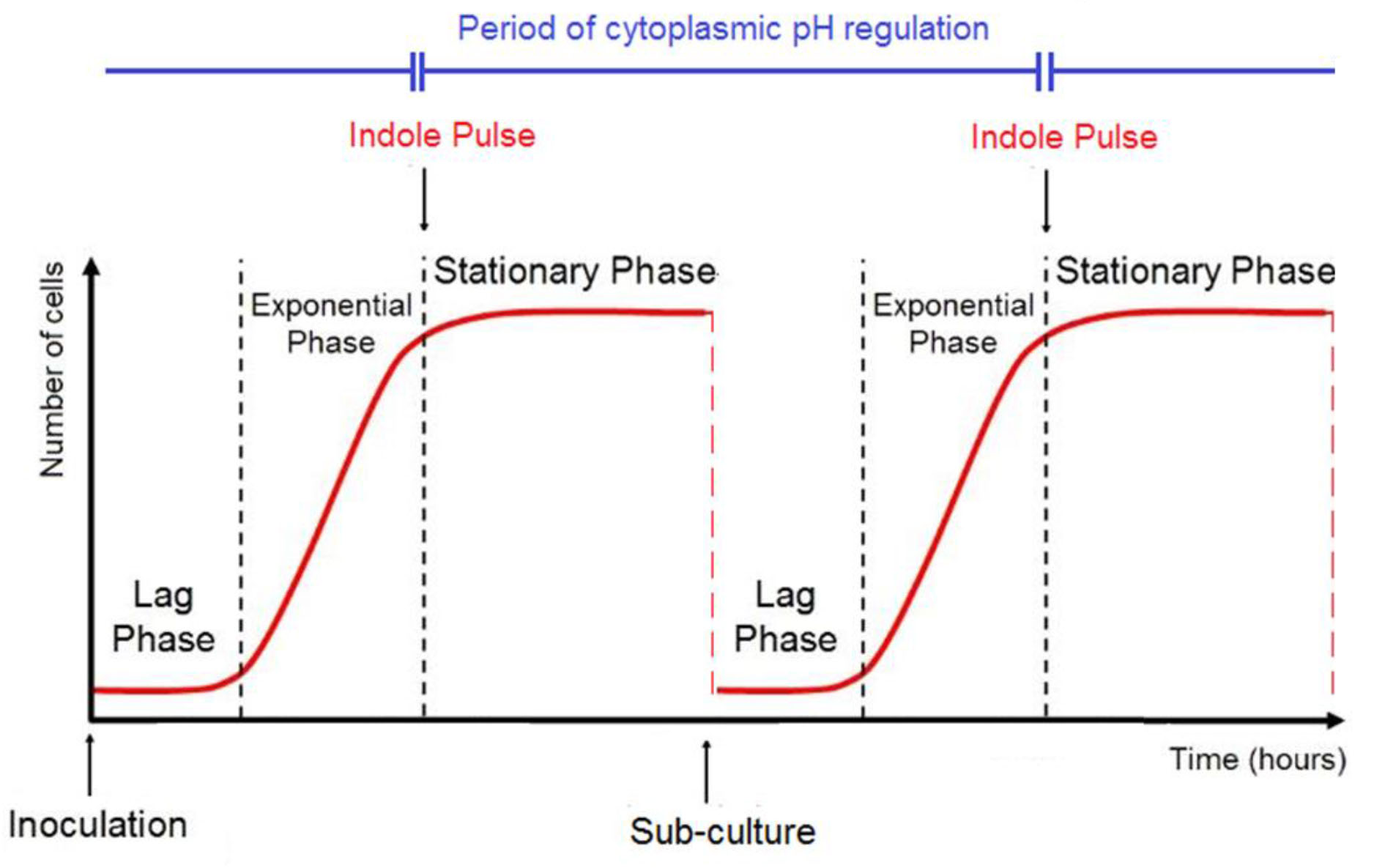
The pH Cycle. A schematic diagram of two consecutive growth cycles for *E. coli*, showing the three main phases of each curve: lag, exponential and stationary phase. The timing of indole pulse in each cycle is marked with downward-pointing arrows, and the period between consecutive pulses over which cytoplasmic pH regulation is maintained is indicated.

The fact that the cytoplasmic pH of wild-type *E. coli* in exponential phase depends upon the growth conditions in a previous overnight culture, rather than the present growth conditions, implies some form of cellular “memory”. In chemosensing *E. coli* uses an external stimulus to change the internal methylation level of chemoreceptors, providing a working memory of the stimulus^33^. *E. coli* has also been reported to retain a memory of the timing of stationary phase entry and to use it to regulate the timing of reinitiation of growth when culture conditions improve. Although the molecular basis of the memory receptor is unknown, it was reported to last for several days without being influenced by stress and environmental changes^34^.

Our experiments suggest that indole causes alteration in the cytoplasmic pH by allowing protons to pass through the cytoplasmic membrane. The main evidence for this is that an artificial pulse with a non-biological ionophore (CCCP or DNP) produces the same outcome as an artificial indole pulse (Fig. 3b). Additionally, the quantitative effect of the indole pulse depends upon the external pH, implying that indole simply facilitates the flow of protons down an existing pH gradient (Fig. 4a and b). It was shown previously that an indole concentration of 3 mM or above is required to reduce the cytoplasmic membrane potential of live *E. coli* cells^5^. This is approximately ten-fold greater than the concentration of indole in the supernatant of a stationary phase culture. In the present work we used membrane vesicles to determine the concentration of indole required to equilibrate the pH across a membrane and found that this occurred rapidly only when the indole concentration was 2.5 or 5 mM; well above the concentration of indole that is available for persistent signalling (Fig. 4c). Thus, the vesicle experiment reinforces the evidence that indole regulation of cytoplasmic pH requires concentrations characteristic of pulse signalling, and that the key property of indole in this context is its proton ionophore activity.

The effect of the pulse remains long after the peak indole concentration has passed but the mechanism of the pulse memory is unknown. One structure that might possibly retain the memory of the pulse is the cell membrane. *E. coli* cells have two membranes offering distinct permeability barriers and separated by a cell wall. The outer membrane acts as filter for large molecules, with transport controlled *via* transmembrane protein pores (porins), which have a preference for small, hydrophilic molecules. In contrast, the inner membrane lacks porins and has a strong preference for lipophilic molecules, which can simply diffuse across the phospholipid bilayer ^35,36,37,38^. The inner membrane seems the better candidate for retaining a memory of the pulse. One potential store of the pulse memory might be in changes to the membrane structure or to the phospholipid composition, which is known to be involved in environmental adaptation and long-term survival^39^. Alterations in phospholipid composition have recently been shown to impact cellular function and stress adaptation^40^. However, there is no obvious way in which indole can alter the phospholipid composition of the inner membrane. A more plausible hypothesis might be that the high concentration of indole during the pulse triggers some structural alteration of the membrane, altering its permeability to protons or the function of other protein ion channels^41^. Changes in the lipid composition of the membrane during entry into the next stationary phase^42^ might necessitate a further pulse to restore the correct proton permeability.

Membrane proteins are another potential retainer of the pulse memory. For example, the oscillation of the Min proteins was shown to be deactivated by concentrations of indole characteristic of the pulse, thereby preventing formation of the FtsZ ring that is prerequisite for division^5^. A functional relationship has recently been established between the Min system and metabolic enzymes, suggesting that the Min system could alter the membrane location of proteins to modulate their enzymatic activity^43^. Thus, the indole pulse could influence the distribution of one or more membrane proteins, with the new, indole-dependent distribution serving as the memory repository.

Alkaline environments are stressful for bacteria^8^, and *E. coli* responds to alkali with SOS and heat shock responses^44,45^. Cytoplasmic pH regulation in *E. coli* is also important in the response to oxidative and acid stress^46,47^. A recent study proposed that some bactericidal antibiotics kill *via* an increase in cytoplasmic pH and that ionophores can rescue cells from antibiotic killing by facilitating an influx of protons to counteract the rise in cytoplasmic pH^7^. Since an indole pulse decreases the cytoplasmic pH, it seems plausible that it might also afford protection against antibiotic and other stresses. Interfering with the regulation of bacterial cytoplasmic pH might therefore provide a route to enhance killing ability of some antibiotics, and this could be exploited in the fight against the continuing rise in antibiotic resistance and the lack of new antibiotics^48^.

Finally, it is relevant to ask why it is desirable for *E. coli* to alter its cytoplasmic pH in response to the presence of tryptophan. In the mammalian gut there appears to be sufficient free tryptophan to ensure the generation of surprisingly large amounts of indole; concentrations up to 3 or 4 mM have been reported^49^. Under these conditions the lower pH characteristic of growth in LB would presumably be maintained. The loss of free tryptophan and indole production might be indicative of a change to a more stressful environment such as starvation of the host or discharge of the bacterium from the intestine into the external environment. Under these conditions the shift towards a higher pH might possibly be adaptive. Further work would be needed to test the validity of this proposal.

In conclusion, this work demonstrates a new role for indole in regulating the cytoplasmic pH of *E. coli*, increasing the importance of indole signalling as a regulator of bacterial physiology. We have shown that the ionophore activity of indole is responsible for setting the cytoplasmic pH during the pulse and that, once set, the correct pH is maintained in a memory-like manner until the next pulse. The physical basis of this memory remains to be identified in future studies.

## EXPERIMENTAL PROCEDURES

### Chemicals and antibiotics

All chemicals and antibiotics were purchased from Sigma. Minimal medium (M9) was made as follows: 1x M9 salts, CaCl_2_ (0.1 mM), MgSO_4_ (1 mM) and FeCl_3_ (0.06 mM) in distilled water. Medium was autoclaved before addition of 0.4 % of glucose (sterile). Supplemented M9 was made by adding the following components (sterile): 0.2 % casamino acids, 1 mg/L thiamine hydrochloride (vitamin B1) and trace elements, including ZnSO_4_ (6.3 µM), CuCl_2_ (7.0 µM), MnSO_4_ (7.1 µM) and CoCl_2_ (7.6 µM). Supplemented M9 + tryptophan was made by adding 1 mM tryptophan (sterile) to the previous recipe.

### Construction of mCherry-pHluorin

A plasmid containing super-ecliptic pHluorin^15,27^ (GenBank AY533296) was a kind gift of Adam Cohen (Department of Physics, Harvard University, USA). A plasmid containing mCherry^28^ (GenBank AY678264) was a kind gift of Marco Geymonat (Department of Genetics, Cambridge University, UK). PCR was used to amplify mCherry and pHluorin with flanking regions to create an mCherry-pHluorin translational fusion linking the gene of mCherry to the gene of pHluorin by a (Gly-Gly-Ser) x2 bridge to give some flexibility to the complex^29^. A pBAD-TOPO vector (Invitrogen) was amplified by PCR and both mCherry and pHluorin fragments were ligated into the vector fragment using a Gibson assembly^30^ kit (NEB) to generate a pBAD TOPO plasmid containing an mCherry-pHluorin translational fusion, pSCM001. The sequence of primers used to generate the pSCM001 plasmid are available in supplementary materials (Supplementary Table 1).

### Strains and culture conditions

The fusion construct, pSCM001, was transformed into *E. coli* BW25112 and BW25113 Δ*tnaA* (kanamcyin resistant: Km^R^), which were both obtained from the Keio collection^31,32^. Cells were cultured routinely at 37°C, with shaking at 120 rpm in Luria Bertani (LB) medium containing 100 µg/ml carbenicillin (carb) to maintain the plasmid. Cultures were streaked to single colonies on LB-carb agar plates and incubated overnight at 37°C to generate stock plates. An independent colony was picked from the stock plate to inoculate LB-carb broth containing 5 mg/L arabinose, Trizma-buffered LB-carb broth (50 mM Trizma, pH 7.0 or 8.0) containing 5 mg/L arabinose or M9-carb (supplemented when required) containing 120 mg/L arabinose and incubated at 37 °C in the shaking incubator. The inducer concentration for LB, Trizma-buffered LB and M9 was determined as the highest concentration that had minimal effect on bacterial growth. Overnight cultures were diluted to OD_600_ = 0.05 in a fresh LB containing 5 mg/L arabinose, fresh Trizma-buffered LB (50 mM Trizma, pH 7.0 or 8.0) containing 5 mg/L arabinose or fresh M9 (supplemented when required) containing 120 mg/L arabinose and allowed to grow to OD_600_ = 0.1 before samples were removed for subsequent assays.

Where required, indole (dissolved in ethanol) was added at appropriate concentrations (30 µM, 50 µM, 100 µM, 250 µM and 500 µM) to BW25113 Δ*tnaA*; ethanol alone was added to controls where appropriate. To impose an artificial pulse, indole (5 mM), CCCP (100 µM) or DNP (800 µM) (all dissolved in ethanol) was added to BW25113 Δ*tnaA* at OD_600_ = 0.1; ethanol alone was added to controls where appropriate. Cells were incubated for 20 min at 37 °C in the shaking incubator and were harvested by centrifugation at 2755 x g for 10 min (Eppendorf 5810 R centrifuge). The supernatant was removed and the cells were resuspended in an equal volume of PBS (pH 7.0). Traces of Indole, CCCP or DNP were removed by centrifugation and removal of the supernatant. Cells were resuspended in an equal volume of fresh LB containing 5 mg/L arabinose and allowed to grow (37 °C, shaking incubator) to OD_600_ = 0.1 before samples were removed for pH measurement.

### Cytoplasmic pH measurement by culture fluorescence

Fluorescence spectroscopy (Agilent Cary Eclipse Fluorescence Spectrophotometer, Santa Clara, CA, USA) was used to measure the fluorescence emission intensity of pHluorin (Ex/Em = 488/510 nm) and mCherry (Ex/Em = 587/610 nm). Sampling was started from OD_600_ = 0.1 for a period of ~18 hrs. For each sample, in addition to measuring the intensity of pHluorin and mCherry in the fluorimeter, the pH of the culture medium was measured using a pH meter. The cytoplasmic pH was calculated from the intensity ratio of pHluorin/mCherry using a standard curve, which was generated as follows. Four additional samples were taken at OD_600_ = 0.6 and the culture medium pH was adjusted manually to 6.5, 7.0, 7.5 and 8.0, followed by addition of 250 µM CCCP, which results in a cytoplasmic pH similar to the pH of the media. Cells were left for 20 min before measuring the intensity of pHluorin and mCherry in the fluorimeter. A new standard curve was generated when the strain and/or the medium was changed.

### Cytoplasmic pH measurement by flow cytometry

Samples with OD_600_ > 0.1 were diluted in ice-cold PBS buffer (pH 7.0) to OD_600_ ≈ 0.1. Samples for confirming the effect of pH on pHluorin (and the lack of any effect on mCherry) were taken at OD_600_ = 0.6 and the external pH was manually adjusted to 6.5, 7.0, 7.5 or 8.0 followed by a 20 min treatment with 250 µM CCCP to equilibrate internal and external pH before diluting in ice-cold PBS buffer (pH 7.0) to OD_600_ ≈ 0.1. All samples were kept on ice before being analysed in a Cytek DxP8 FACScan (Cytek Bioscience, Fremont, CA, USA). Fluorescence was excited at 488 nm for pHluorin and 561 nm for mCherry and measured through an emission filter of 530 nm and 615 nm, respectively, with 25 nm width. For each sample, 50,000 events were collected at a rate between 1000 and 2000 events per second. Data were collected and analysed using FlowJo V10 (FlowJo LLC, Ashland, Oregon, USA).

### Intravesicular pH measurements

Large unilamellar vesicles (LUVs) with a diameter of ~ 200 nm were prepared from 2-oleoyl-1-palmitoyl-*sn*-glycero-3-phosphocholine (POPC) with encapsulating HPTS (POPC‐LUVs⊃HPTS). 1 ml 1x PBS (pH 7.4) containing 10 µl POPC‐LUVs⊃ 1mM HPTS (pH 7.4) was used for each experiment. 5 µL indole in ethanol was added to mix with the LUVs at t = 1.5 min, and then a pH gradient (ΔpH=0.9) across the vesicle membrane was generated by adding 4 µl of 10% HCl to the extravesicular buffer at t = 2 min, lowering the external pH value to 6.5. Therefore, solution inside POPC-LUV after HCl addition: 1 mM HPTS, 1x PBS, pH 7.4; solution outside POPC-LUV after HCl addition: 1x PBS, pH 6.5. The fluorescence intensity of HPTS was monitored for ~ 6 min before lysing POPC-LUV by the addition of 20 µl of 10% Triton X-100 to obtain a final normalisation point (HPTS intensity corresponds to pH = 6.5). The fluorescence intensity of HPTS (*λ*_ex_=450 nm, *λ*_em_=510 nm) in POPC‐LUVs⊃HPTS throughout the experiment was measured by fluorescence spectroscopy (Agilent Cary Eclipse Fluorescence Spectrophotometer, Santa Clara, CA, USA). The experiment was repeated with the addition of a range of indole concentrations (0.5, 1.0, 2.5 and 5.0 mM) or 5 µL pure ethanol as a blank control (0 mM indole) to the extravesicular buffer before the addition of HCl, and changes in HPTS fluorescence were monitored. Each experiment (with and without indole) was performed in triplicate.

## ACKNOWLEDGEMENTS

This work was supported by funding from the Leverhulme Trust, UK (RPG-2015-184). J.Z. and U.F.K. thank the support from UK Engineering and Physical Sciences Research Council (EPSRC, EP/M008258/1). J.C. acknowledges support from the Biotechnology and Biological Sciences Research Council. K.A.N. was supported by Winton Programme for the Physics of Sustainability, Trinity-Henry Barlow Scholarship, National Physical Laboratory and ERC consolidator grant (DesignerPores 647144). We thank Annabel Murphy, University of Cambridge Department of Chemistry for providing advice on the use of C18 columns.

## AUTHOR CONTRIBUTIONS

A.Z., J.C., U.F.K., and D.K.S. designed the study. S.C-M. constructed the pHluorin-mCherry plasmid. J.Z., and K.A. performed the vesicles experiment. A.Z. performed all other experiments. A.Z., and D.K.S. wrote the manuscript.

